# A pyrethroïd-treated bed net increases host attractiveness for *Anopheles gambiae s.s.* carrying the *kdr* allele in a dual-choice olfactometer

**DOI:** 10.1101/077552

**Authors:** Angélique Porciani, Malal Diop, Nicolas Moiroux, Tatiana Kadoke-Lambi, Anna Cohuet, Fabrice Chandre, Laurent Dormont, Cédric Pennetier

## Abstract

The use of long lasting insecticide nets (LLINs) treated with pyrethroïd is known for its major contribution in malaria control. However, LLINs are suspected to induce behavioral changes in malaria vectors, which may in turn drastically affect their efficacy against *Plasmodium sp*. transmission. In sub Saharan Africa, where malaria imposes the heaviest burden, the main malaria vectors are widely resistant to pyrethroïds, the insecticide family used on LLINs, which also threatens LLIN efficiency. There is therefore a crucial need for deciphering how insecticide-impregnated materials might affect the host-seeking behavior of malaria vectors in regards to insecticide resistance. In this study, we explored the impact of permethrin-impregnated net on the host attractiveness for *Anopheles gambiae* mosquitoes, either susceptible to insecticides, or carrying the insecticide resistance conferring allele *kdr*. Groups of female mosquitoes were released in a dual-choice olfactometer and their movements towards an attractive odor source (a rabbit) protected by insecticide-treated (ITN) or untreated nets (UTN) were monitored. *Kdr* homozygous mosquitoes, resistant to insecticides, were more attracted by a host behind an ITN than an UTN, while the presence of insecticide on the net did not affect the choice of susceptible mosquitoes. These results suggest that permethrin-impregnated net is detectable by malaria vectors and that the *kdr* mutation impacts their response to a LLIN protected host. We discuss the implication of these results for malaria vector control.

## Introduction

*Anopheles gambiae* is one of the major mosquito vectors of human malaria parasites in sub-Saharan Africa. Its remarkable vectorial capacity [1] mainly relies on its high degree of anthropophily. Moreover, *An. gambiae* prefers to bite humans indoors and often rests inside houses after blood feeding [2–4]. These behavioral preferences led to the development of insecticide-based indoor vector control measures, such as insecticide-treated bed nets (ITNs) and indoor residual spraying (IRS), to limit the human-vector contacts and reduce mosquito survival. To date, four insecticide families are available for IRS (organochlorides, organophosphates, carbamates and pyrethroïds), whereas only pyrethroïds are recommended for mosquito nets because of their low mammalian toxicity and high insecticidal potency [5].

To kill, insecticide molecules must contact and penetrate through the mosquito cuticle/gut to then reach and interact with their target before being degraded. Any physiological or behavioral mechanism that may interfere with one of these steps can lead to insecticide resistance. The widespread use of pyrethroïd (PYR) insecticides in malaria vector control and agriculture has favored the development of resistance in malaria vector species [6]. One of the most studied physiological mechanisms involved in PYR resistance is the reduced sensitivity of the voltage-gated sodium channels to PYR binding caused by non-silent mutations, known as knockdown resistance (*kdr*) mutations [7]. Behavioral resistance is another mechanism involved in PYR resistance. This can be defined as a modification of the mosquito behavior to avoid contact with a lethal dose of insecticide [8]. To date, behavioral resistance to insecticides remains poorly documented, despite of its huge potential impact on malaria transmission.

Behavioral adaptations to pesticides can be classified as stimulus-dependent or -independent [9]. Stimulus-independent adaptations are not associated with the perception of chemicals, but more probably with modifications of the vector intrinsic behavior, such as changes in host-seeking behavior preferences (level of anthropophily, endophagy, endophily or hourly biting activities). Such behavioral modifications have recently been observed in the context of ITN widespread use: mosquito vectors may postpone their bloodfeeding until the morning, when human hosts are protected by ITNs anymore [10–12]. These changes may limit the contact between aggressive malaria vectors and treated surfaces, thus threatening the efficiency of indoor vector control tools. Conversely, stimulus-dependent behavioral adaptations are specifically linked to the detection of chemicals. Stimulus-dependent insecticide avoidance can be defined as a “fly away” behavior to leave the immediate toxic environment after contact (irritancy) or not (repellence) with the treated surface [13–15]. Avoidance behavior following contact with PYR has been reported in some cases [16–20], but similar behavior in the absence of direct contact with the insecticide has been poorly documented. Only indirect observations suggest a detection and avoidance of ITNs by malaria vectors: mosquito entry rates were found reduced in experimental huts containing insecticide-treated nets compared to entry rates in control huts, moreover the observed rates were dependent on the *kdr* allele presence in the mosquitoes [21–23]. Although the effects of pyrethroïds on different part of host seeking behavior has been already studied [20,24–26], their influence on the relative host attractiveness has been neglected despite its importance in host choice and on malaria transmission. Therefore, in order to adequately evaluate and use ITNs, it has become urgent to investigate the possible modulation of the host-seeking behavior in presence of indoor vector control tools in regards to other insecticide resistance mechanisms

In this study, we examined the long-range host-seeking behavior of *An. gambiae* mosquitoes to determine whether the attractiveness of a vertebrate host (a rabbit) in a dual-choice olfactometer was influenced by physical and/or chemical barriers (insecticide-treated and untreated nets) and by the mosquito *kdr* (L1014F) genotype.

## Methods

### Ethics statement

Rabbits were handled and blood drawn in accordance to the protocol approved by National Comity for Ethic and Research (CNERS) and Health ministry of Benin (N°023). This study was carried out in strict accordance with the recommendations of Animal Care and Use Committee named “Comité d’éthique pour l’expérimentation animale; Languedoc Roussillon” and the protocol was approved by the Committee on the Ethics of Animal Experiments (CEEA-LR-13002 for the rabbits). Rabbits were not subjected to anesthesia, analgesia or sacrifice.

### Mosquitoes

Two laboratory reference strains of *Anopheles gambiae sensu stricto* (formerly called S molecular form) (20) were used in this study. The Kisumu reference strain, isolated in Kenya in 1975 (VectorBase, http://www.vectorbase.org, KISUMU1), is free of any detectable insecticide resistance mechanism. The *kdr*-kis strain was obtained by introgression into the Kisumu genome of the *kdr*-west allele (L1014F) [27] that originated from a PYR-resistant population collected in Kou Valley, Burkina Faso, which was used to establish a strain named VKPer. Introgression was obtained through 19 successive back-crosses between Kisumu and VKPer [28]. VKPer strain displayed the same expression level of metabolic resistance enzyme as Kisumu [29]. Kisumu and *kdr*-kis mosquitoes are therefore homozygous susceptible (SS) and homozygous resistant (RR) at the *kdr* locus, respectively. The heterozygous genotype RS was obtained by crossing Kisumu SS females with *kdr*-kis RR males.

Mosquitoes were reared in insectary conditions (27±3°C, 60-80% relative humidity and a 12:12 light and dark cycle). Ground cat food was used to feed larvae and 10% sucrose solution (with rabbit blood twice per week) to feed adult females. For behavioral experiments, 5-12 day old females, without prior access to a blood meal, were starved for 4h before the assay.

### Experimental set-up

The dual-choice olfactometer was adapted from Geier and Boeckh (1999) [30]. It was made of Plexiglas and was divided in four parts: release zone (RZ), flight chamber (FC) and one collecting zone in each of the two arms (A1 or A2) (Fig 1). Rotating doors made from mesh gauze in the RZ and in both arms allowed mosquito release or capture. The upwind part of the experimental set-up was composed of a wide chamber where an attractive host (a rabbit) can be placed, and that was connected to two treatment boxes that contained or not the nets. Each treatment box was connected to one arm of the olfactometer. In order to avoid any perturbation on the airflow by the treatment, fans were placed on the downwind faces of the experiment boxes and extracted the air from the treatment boxes to the olfactometer, providing the odor-laden air current. At the beginning of each experiment, the airflow was measured in arm 1 and 2 and in the release zone using a Testo^©^ 435-1 multifunctional meter (Testo, Forbach, France) and thermo-anemometric probe (m.s^−1^) and adjusted at 0.20 ± 0.03 m.s^−1^. During the experiment, a thick black tarpaulin covered the olfactometer to keep all the system in darkness and avoid visual disturbance.

**Fig 1.**
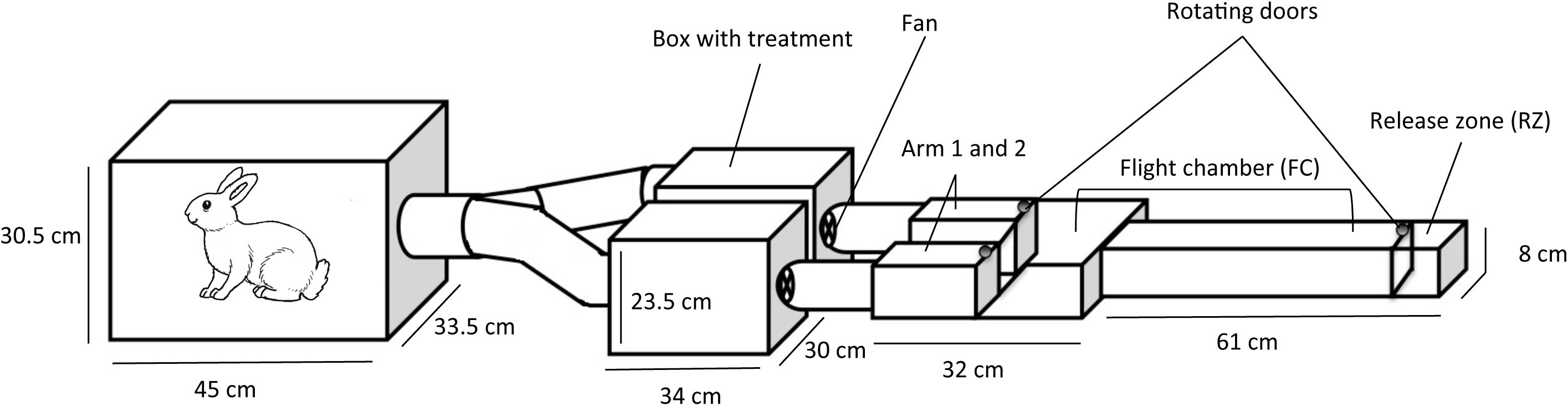
Experimental set-up. Dual-choice olfactometer (right side) connected to the treatment boxes (middle) and the wide chamber (left side).

### Experimental design

Four experiments, summarized in Table 1, were performed using SS, RR and RS mosquitoes. The treatment boxes and the wide chamber were empty during the first experiment. For the other experiments, the wide chamber contained a rabbit as odor source. The treatment boxes contained, depending on the experiment, nothing or 2m^2^ of untreated (UTN) or insecticide-treated net (ITN, Olyset^®^ Net impregnated with 1000mg/m^2^ of permethrin). Nets were divided in 50 pieces of 20x20cm and hung on a metallic structure perpendicularly to the air flow. The same nets were used during all experiment, the Olyset^®^ was conserved at 4°C between each day of experiment. The nets were placed in boxes that could not be visible for mosquitoes, so that no visual clues were available to mosquitoes during the experiments.

**Table 1:** Description of the experimental design

| Experiment no. | Experiment name | Odor source | Treatment box 1 | Treatment box 2 |
| --- | --- | --- | --- | --- |
| 1 | Empty | None | Empty | Empty |
| 2 | Rabbit alone | Rabbit | Empty | Empty |
| 3 | Rabbit + UTN | Rabbit | Empty | UTN |
|  |  |  | UTN | Empty |
| 4 | Rabbit + ITN | Rabbit | ITN | UTN |
|  |  |  | UTN | ITN |

(UTN: untreated net, ITN: insecticide-treated net)

Assays for the four experiments were performed every day for 20 days between 10:00am and 14:00pm (corresponding to mosquito strain feeding time in laboratory). We always started with assays of experiment 1, to check possible odor or insecticide contaminations. When possible (i.e., when the insectary production was sufficient), females of the three genotypes were tested the same day for the four experiments, otherwise at least two genotypes were tested the same day (a summary of the assays is presented in supplementary data). Each day, in assays for experiments 3 and 4, treatments were rotated one time between boxes to prevent any arm effect. Between rotations, the boxes were carefully cleaned with ethanol to avoid any residual insecticide effect.

Moreover, the olfactometer was cleaned with ethanol every day. The experimenter wore latex gloves to avoid contamination. The same rabbit was used as odor source during all the study. It was a 1-year old female reared in the same conditions as those used in insectaries to feed mosquitoes. CO_2_ concentration and relative humidity (RH) were monitored in each arms with a Testo^©^ 435-1 multifunctional meter (Testo, Forbach, France) equipped with an Indoor Air Quality (IAQ) probe [%RH; range: 0 to +100 %RH; accuracy: ±2 %RH (+2 to +98 %RH)], [CO_2_; range: 0 to +10000 ppm_;_ accuracy: (±75 ppm CO_2_ ±3% of mv) (0 to +5000 ppm CO_2_)]. The room was kept at a constant temperature of 25°C during the study.

For each assay, a batch of 20-23 females was released in the RZ for 5 min for acclimation. The rotating doors were then opened and females were free to fly in the olfactometer. After 5 minutes, the rotating doors were closed and the numbers of mosquitoes in RZ (N_RZ_), FC (N_FC_), A1 and A2 (N_A1_ and N_A2_) were recorded (Figure 1).

### Behavioral indicators

The indicators used in this study describe the mosquito progress inside the olfactometer and the relative attractiveness (RA) of treatments or arms.

Two indicators of the progression inside the olfactometer were calculated. First, upwind flight (UF) that is the proportion of female that left the release zone (i.e. collected in FC, A1 and A2) relative to the total number of released mosquitoes (N). Second is the localization (L) of odor source that is the proportion of female that reached A1 and A2 (N_A1_ and N_A2_), relative to the number of mosquitoes that left the RZ (N - N_RZ_). These indicators were calculated for each release and for each odor source (none, rabbit without ITN and rabbit with ITN).

The upwind flight and localization values measured in experiment 1 (empty set-up, clean air) are baseline indicators of the anemotactic response of the three mosquito genotypes to air flow. The influence of rabbit odor on mosquito’s progression was determined by comparing the values of upwind flight and localization recorded in the empty system (experiment 1) with those recorded in the system without ITN (merged UF and L values of experiments 2 and 3). The merged upwind flight and localization values recorded in experiments 2 and 3 (rabbit odor, no ITN) were compared to those recorded in experiment 4 (rabbit odor and ITN) to determine ITN odor influence on mosquito behavior.

The relative attractiveness (RA) of one arm versus the other was calculated as the proportion of mosquitoes in A1 or A2 (N_A1_ or N_A2_) relative to the sum of the mosquitoes collected in both arms. In order to verify the symmetry of the experimental set-up, we measured RA_exp2_ in experiment 2 (rabbit as an odor source, empty boxes) as follow and expected it to not be different than 0.5: 
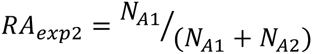

Relative attractiveness of UTN versus empty box (RA _exp3_) and ITN versus UTN (RA _exp4_) were also calculated from experiments 3 and 4, respectively, using the following equations: 
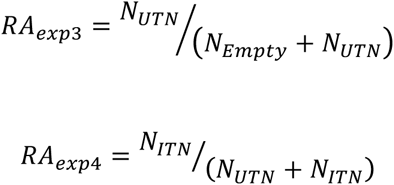
where *N*_*UTN*_ is the number of mosquitoes collected in the arm with the box containing the UTN (experiment 3 or 4), *N*_*Empty*_ is the number of mosquitoes in the arm with the empty box (experiment 3) and *N*_*ITN*_ is the number of mosquitoes collected in the arm with the box containing the ITN (experiment 4). The measure of RA _exp3_ allowed us to assess the possible effect of the UTN as a physical barrier for the diffusion of odor coming from the rabbit to the olfactometer.

### Statistical analysis

All analyses were performed using the R software, version 3.0.2 [31], with the lme4 package [32]. We analyzed upwind flight and localization using binomial logistic mixed-effect models. The day of release was set as random intercept because releases performed on a same day might not be independent and because all three genotypes have not been tested each day. The *kdr* genotypes (SS, RS or RR), the different odor sources (none, Rabbit without ITN, and Rabbit+ITN) and interactions between them were included in the models as explanatory variables. Upwind flight (UF) and localization (L) models were written as follow: 
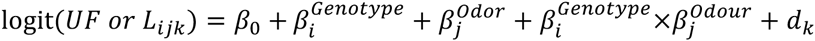
where *UF or L*_*ijk*_ is the proportion UF or L recorded for genotype *i* with odor source *j* on day *k*, 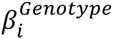 denotes the effect on the logit of the classification in category *i* (SS, RS or RR) of Genotype; 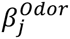denotes the effect of the classification in category *j* of Odor source: Empty (experiment 1), Rabbit without ITN (experiment 2 and 3), or Rabbit+ITN (experiment 4); and d_*k*_ represents the random intercept for day *k*. Each combination of categories *i* and *j* of the explanatory variables was successively used as reference class for multiple comparisons among genotypes and odor sources. Odds ratios and their 95% confidence intervals (CI) were computed.

We verified the symmetry of the experimental set-up by modelling the relative attractiveness measured in experiment 2 (*RA_exp2_*) using a binomial mixed-effect model with the release day as random effect: 
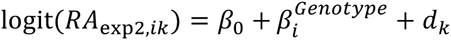
where *RA*_*exp2,ik*_ is the proportion RA in A1 for genotype *i* in experiment 2 on day *k*, 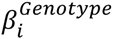is the effect on the logit of the classification in category *i* (SS, RS or RR) of Genotype; and d_*k*_, the random intercept for day *k*.

Relative attractiveness of UTN vs. empty box and ITN vs. UTN were analyzed using a similar model that, in addition, allowed for random effects of the box that received the treatment: 
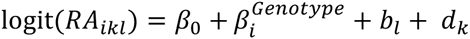
where *RA*_*ikl*_ is the proportion *RA*_*exp3*_ or *RA*_*exp4*_ for genotype *i* on day *k* with the treatment placed in box *l*, 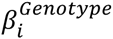indicates the effect on the logit of the classification in category *i* (SS, RS or RR) of Genotype; *b*_*l*_, the effect on the logit of the the box *l* that received the treatment (UTN or ITN for *RA_exp3_* and *RA_exp4_*, respectively) and d_*k*_, the random intercept for day *k*. Each genotype was successively used as reference class for multiple comparisons. Odds ratios and their 95% CI were computed.

CO_2_ concentrations were compared between arms using the Wilcoxon signed-rank test for paired data. RH values were compared between arms using the paired T test.

## Results

Overall, 6286 mosquitoes were included in the assays (2621 SS, 1268 RS and 2397 RR) during 47, 49, 84 and 98 releases for experiments 1 to 4 respectively (Table 2).

**Table 2:**
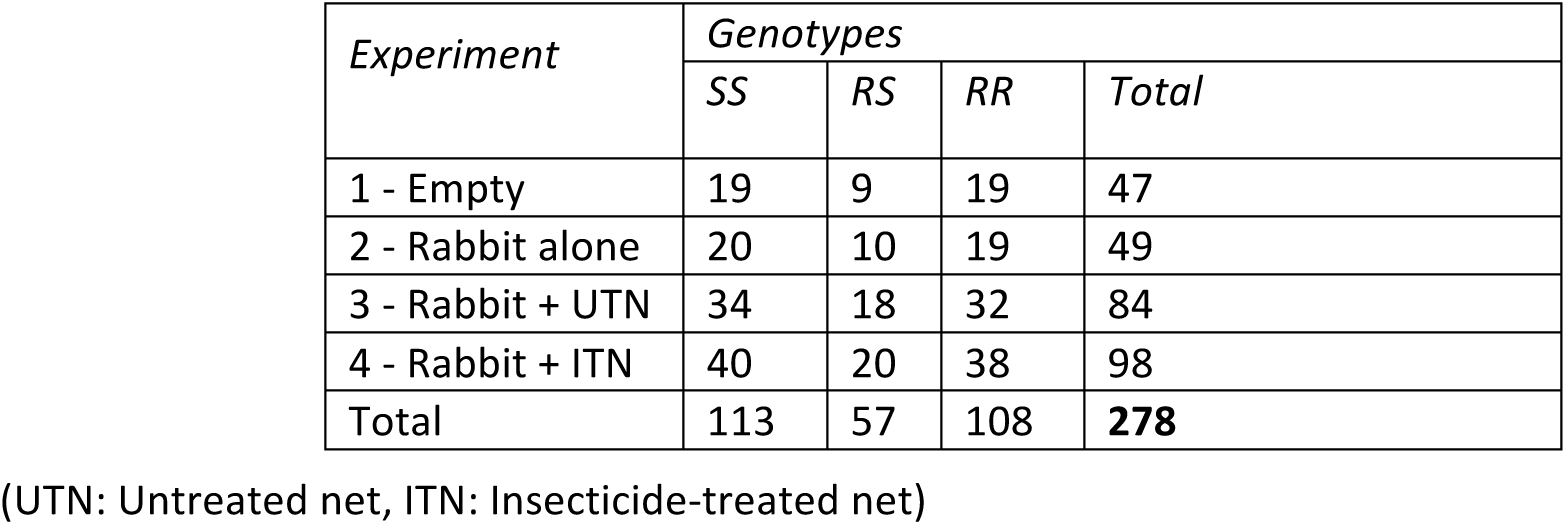
Number of releases performed per genotype and experiment

### Do *An. gambiae* females respond to the air flow?

We first investigated the response to the airflow (anemotactic response) by calculating the proportion of upwind flight (UF) females and those located (L) in arms in the empty set-up (Experiment 1). Overall, the probability to leave RZ (UF) was 0.43 (95%CI [0.38 – 0.48]; Fig 2A). Among the activated mosquitoes, 10% (95%CI [6 – 17]) reached A1 or A2 (Fig 2B). In spite of similar upwind flight proportion among genotypes, the probability of localization (L) for RS anopheles was higher than those of RR mosquitoes (Figure 2B; OR_L_= 2.15, 95%CI [1.04, 4.41]).

**Fig 2:**
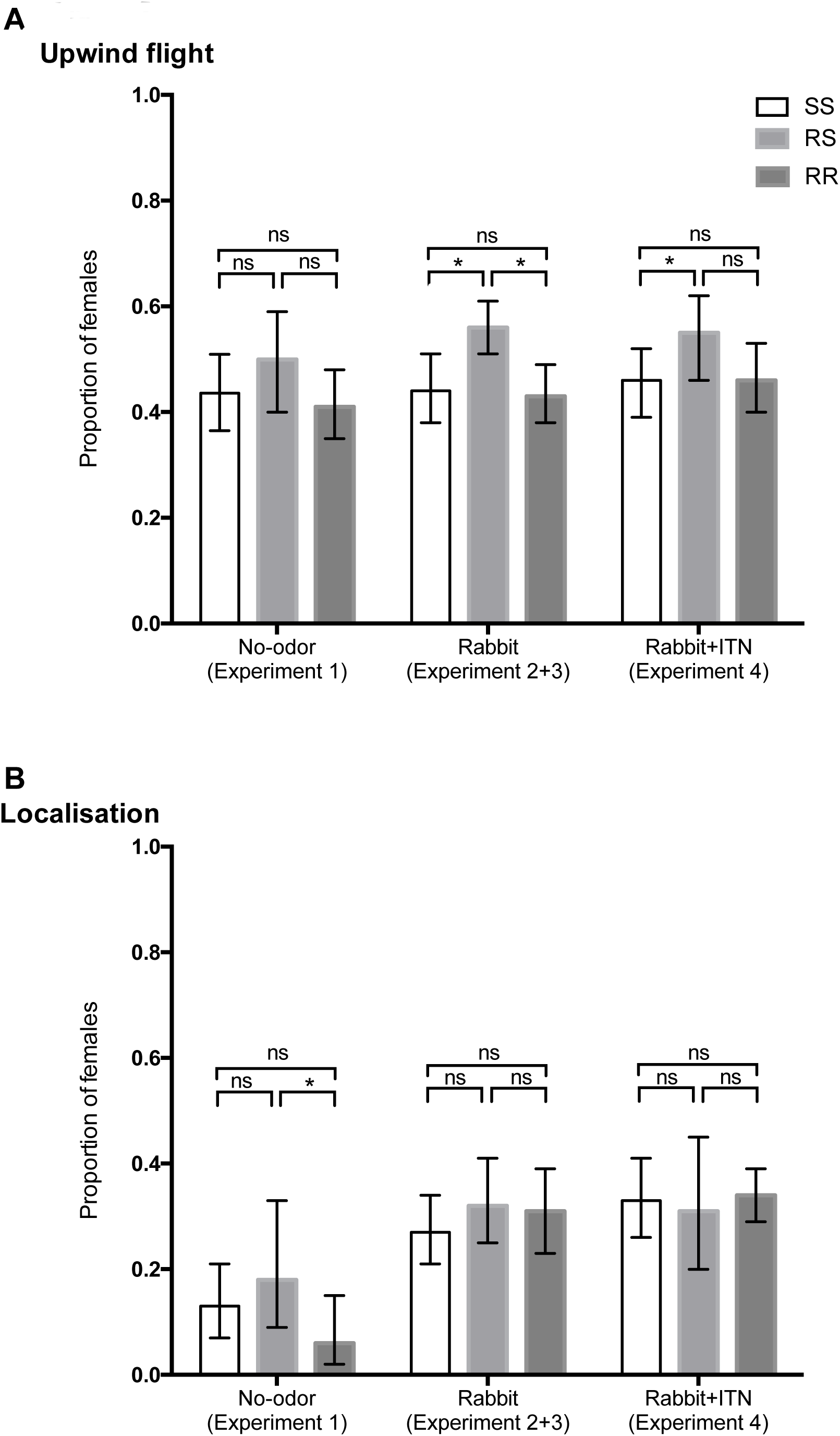
Upwind flight and localization indicators for the three genotypes in relation to treatment. (Mean±95% Confidence Interval). ***p<0.001, **p≤0.01, *p≤0.05, ns= not significant.

### Do *An. gambiae* females respond to an attractive odor source?

The presence of a rabbit as an attractive odor source (experiments 2 and 3) did not change the proportion of upwind flight mosquitoes compared to the experiments without attractant odor (experiment 1), independently of their genotype (Table 3). However, the comparison of the upwind flight probability between genotypes show that for RS mosquitoes, UF probabilities became significantly higher than for SS and RR individuals (Fig 2A; OR_RSvsSS_ = 1.24 95%CI [1.01, 1.54]; OR_RSvsRR_ = 1.29 95%CI[1.04, 1.59]). Moreover, the localization probability significantly increased for all genotypes in the presence of an odor stimulus compared to no odor (Table 3), independently of genotypes (Fig 2B). The rabbit odor had an effect on mosquito behavior only when they were close to arms likely because of the odor concentration that was more important in arms than in the release zone.

**Table 3:**
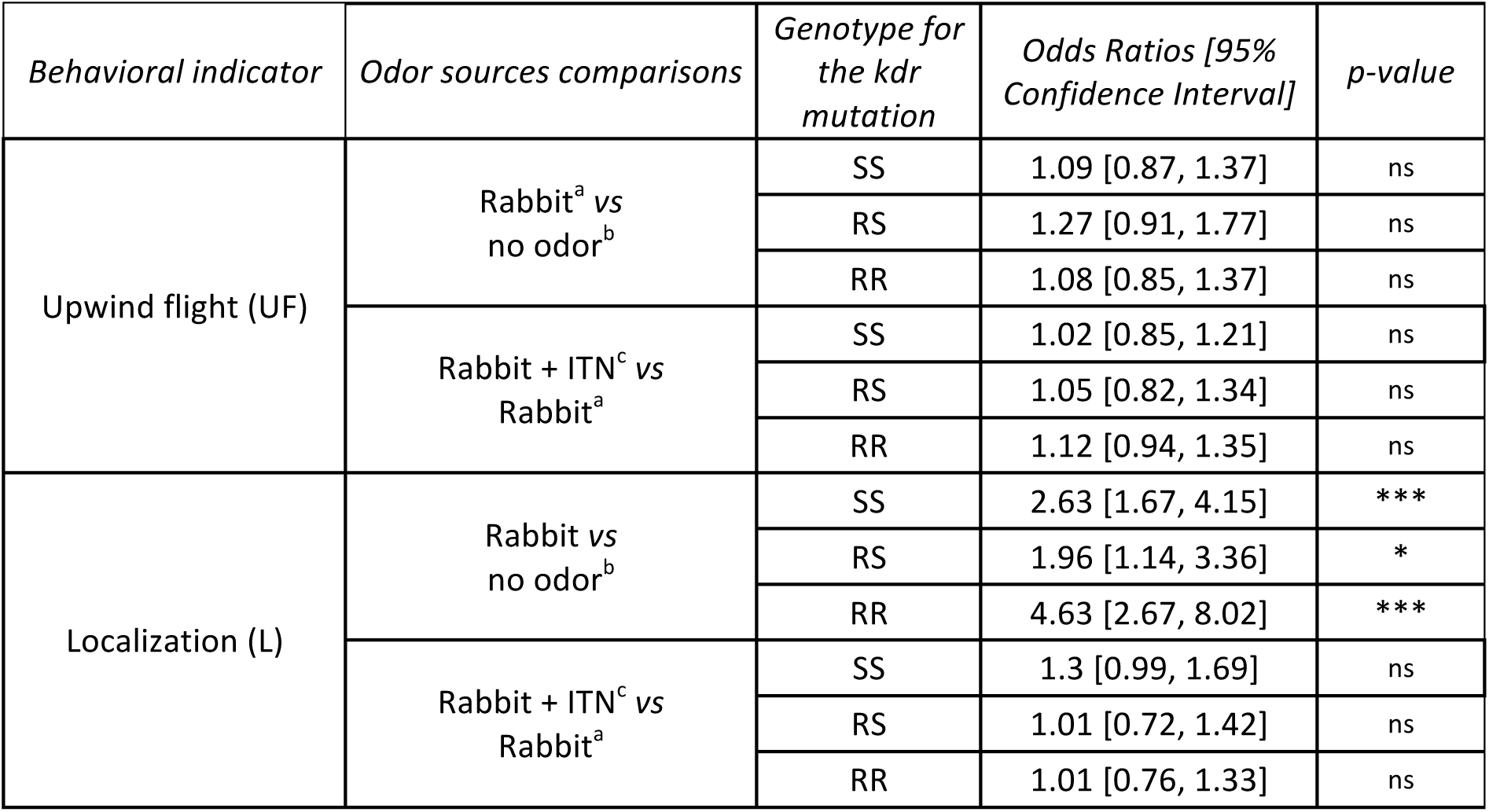
Results of the Upwind flight (UF) and localization (L) generalized linear models. Comparison of mosquitoes’ progress first when the rabbit was added as an odor source (*vs.* no odor) and then when ITN was present (*vs*. rabbit alone). ^a^ experiments 2 and 3, ^b^ experiment 1, ^c^ experiment 4 (see Table 1); ***p <0.001, **p≤ 0.01, *p≤0.05, ns: not significant. ITN: insecticide-treated net. SS: homozygous for the susceptible allele, RS: heterozygous, RR: homozygous for the resistant allele.

| Behavioral indicator | Odor sources comparisons | Genotype for the *kdr* mutation | Odds Ratios [95% Confidence Interval] | p-value |
| --- | --- | --- | --- | --- |
| Upwind flight (UF) | Rabbit<sup>a</sup> vs no odor<sup>b</sup> | SS | 1.09 [0.87, 1.37] | ns |
|  |  | RS | 1.27 [0.91, 1.77] | ns |
|  |  | RR | 1.08 [0.85, 1.37] | ns |
|  | Rabbit + ITN<sup>c</sup> vs Rabbit<sup>a</sup> | SS | 1.02 [0.85, 1.21] | ns |
|  |  | RS | 1.05 [0.82, 1.34] | ns |
|  |  | RR | 1.12 [0.94, 1.35] | ns |
| Localization (L) | Rabbit vs no odor<sup>b</sup> | SS | 2.63 [1.67, 4.15] | *** |
|  |  | RS | 1.96 [1.14, 3.36] | * |
|  |  | RR | 4.63 [2.67, 8.02] | *** |
|  | Rabbit + ITN<sup>c</sup> vs Rabbit<sup>a</sup> | SS | 1.3 [0.99, 1.69] | ns |
|  |  | RS | 1.01 [0.72, 1.42] | ns |
|  |  | RR | 1.01 [0.76, 1.33] | ns |

### Is mosquito response influenced by insecticide-treated nets?

To test whether the insecticide on net fibers affected mosquito progression, we compared upwind flight and localization probabilities in the presence (experiment 4) or absence (experiments 2 and 3) of the ITN. The probabilities were similar in presence or absence of the ITN, regardless of the genotype (Table 3; Fig 2A, 2B). Nevertheless, the comparison between genotypes showed that upwind flight probabilities for heterozygous RS mosquitoes remained higher than those of the two other genotypes, both in the presence or absence of insecticide (Fig 2A; OR_RS*vs*SS_= 1.28 95%CI [1.01, 1.62], OR_RS*vs*RR_= 1.20 95%CI [0.94, 1.53]).

### Is the experimental set up symmetric?

Analysis of the arms’ relative attractiveness data from experiment 2 (Rabbit odor; two empty boxes) showed no significant differences between the number of mosquitoes collected in A1 *vs.* A2, regardless of the genotypes (Fig 3A; RA_exp2,SS_ = 0.58, 95%CI [0.34, 0.79]; RA_exp2,RS_ =0.62, 95%CI [0.34, 0.83]; RA_exp2,RR_ = 0.54, 95%CI [0.30, 0.76]). No difference in RA_exp2_ was observed among genotypes (OR_SS*vs*RS_= 1.16 95%CI [0.46, 2.94]; OR_SS*vs*RR_=0.85 95%CI [0.38, 1.94], OR_RS*vs*RR_= 0.73 95%CI [0.28, 1.90]). Moreover, CO_2_ concentration and RH were not different between arms (p>0.05; S1). These results demonstrated that the olfactometer was symmetrical. Moreover, analyses of RA_exp3_ and RA_exp4_, (results described below), showed no effect relative to the box receiving the treatment (i.e. variable no significant in the model), indicating that symmetry was maintained during experiments 3 and 4.

**Figure 3:**
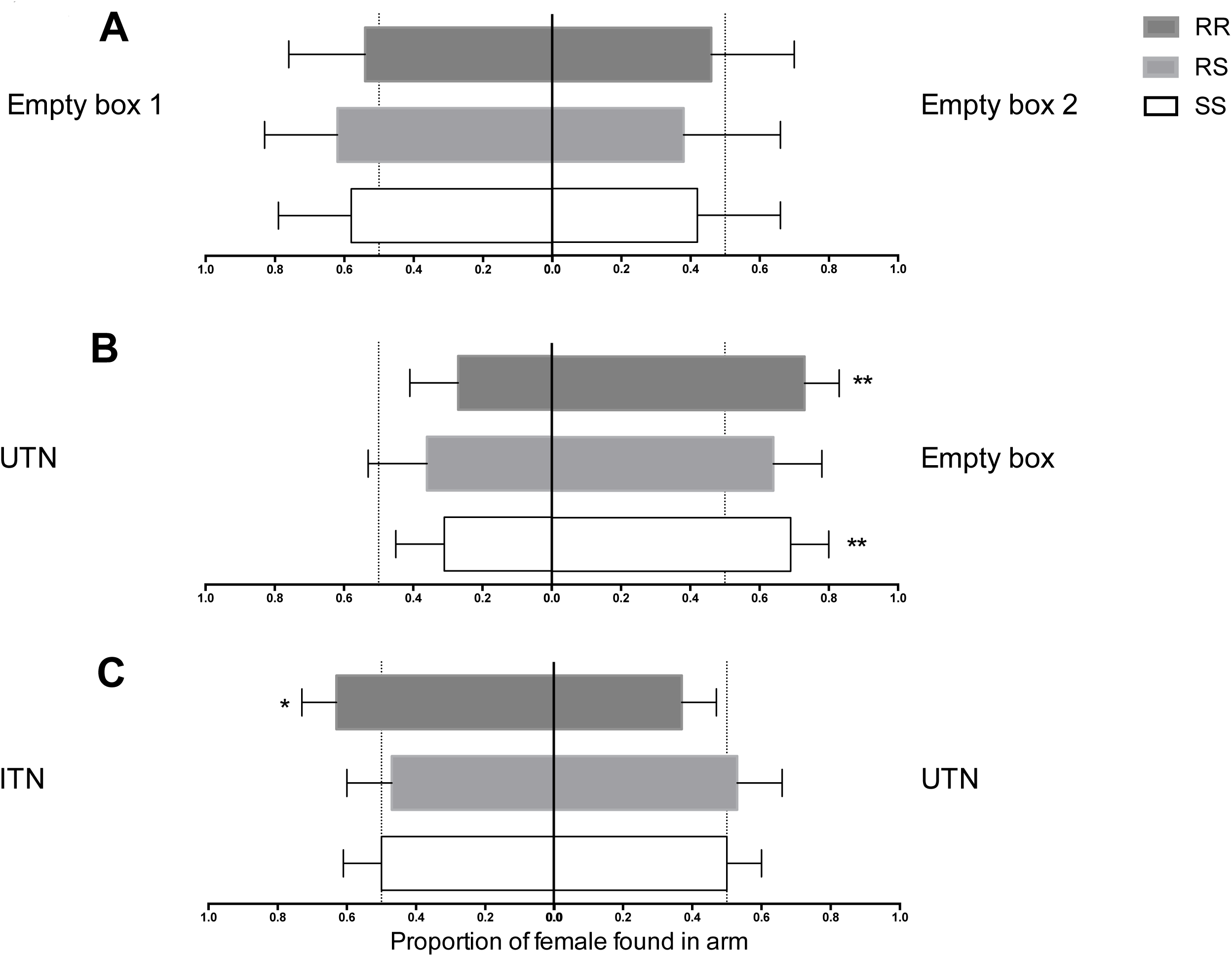
RA: number of mosquitoes found in one arm relative to the total number of mosquitoes found in both arms. (A) Experiment 2 (rabbit only). (B) Experiment 3 (rabbit + UTN or empty box). (C) Experiment 4 (Rabbit+ UTN or ITN). Asterisks show difference to 0.5, traducing a choice for one treatment rather than the other. Error bars show the 95% CI; **p≤0.01, *p≤0.05. UTN: Untreated net, ITN: Insecticide-treated net. SS: homozygote for the L1014S allele (insecticide-susceptible), RS: heterozygous for the L1014F allele, RR: homozygous for L1014F allele (insecticide-resistant).

### Is the attractiveness of the odor source influenced by the UTN?

In experiment 3 (one empty box and one box with 2 m² of UTN, both in presence of rabbit odor), the empty box was more attractive for SS and RR mosquitoes but not for RS (RA_exp3,SS_ = 0,31 95%CI [0.20,0.46], p-value= 0.013; RA_exp3,RR_ = 0.27, 95%CI [0.17,0.42], p-value=0.002)(Fig 3B). No significant difference of mosquito’s proportion in arms was evidenced between genotypes (OR_SS*vs*RS_= 1.24, 95%CI [0.61, 2.50]; OR_SS*vs*RR_=0.83, 95%CI [0.41, 1.66], OR_RS*vs*RR_= 0.67, 95%CI [0.31, 1.47]). CO2 concentration was not different between arms, while a significant 1% difference in RH was observed (63.9 % in the UTN arm and 64.9% in the empty arm, paired T test p-value = 0.007).

### Is the attractiveness of the odor source influenced by the ITN ?

Analysis of RA_exp4_ from experiment 4 (Rabbit odor; one box with 2 m² of UTN and one with 2 m² of ITN) showed that RR mosquitoes preferably chose the ITN arm with probability 0.63 (95%CI [0.53- 0.73], p-value=0.01; Fig 3C). This probability was significantly higher than those observed both for RS (RA_exp4,RS_ = 0.47 95%CI [0.34-0.60]; OR_RR*vs*RS_= 1.95, 95%CI [1.06, 3.57], p-value=0.03) and SS genotypes (RA_exp4,SS_ = 0.5 95%CI [0.40-0.61]; OR_RR*vs*SS_=1.71, 95%CI [1.03, 2.83], p-value=0.04). CO_2_ concentration and RH were not different between arms.

## Discussion

The host-seeking behavior of mosquitoes towards humans sleeping under a bed net is poorly understood. Particularly, it is not known whether specific volatile chemicals emanating from treated nets might modulate this behavior. Here, we used a dual-choice olfactometer to assess whether the presence of permethrin-treated nets may influence the host attractiveness for mosquitoes of different *kdr* genotypes. We found that nets represent both a physical and a chemical signal that modulate mosquito activation and/or choice. Moreover, the three *kdr* genotypes behaved differently in response to host odors in the presence of ITNs or UTNs.

## Physical barrier & environmental cues

In experiment 3, mosquitoes preferably chose the arm connected to the empty box rather than the box with UTNs. No difference in CO_2_ quantity was noted between arms. However, the humidity level was slightly higher in the arm connected with the empty box. As humidity is known to attract mosquitoes [33], the observed preference for the empty box (higher humidity) was not surprising. This difference could have been caused by the physical barrier formed by UTNs that may absorb humidity coming from the rabbit box. In addition, the net structure could also have retained volatile chemicals emanating from rabbit which are important in mosquito orientation and choice [34,35].

## Chemical ecology & ITNs

Our results raised the question of the volatile chemicals emanating from nets that may drive a specific behavior in resistant mosquitoes. Permethrin is not known as a classical volatile compound because of its low vapor pressure (5.18x10^−8^mm Hg at 25°c). Nevertheless, Bouvier et al [41] recently detected permethrin in indoor air samples (11 and 18.8 ng/m^3^ for *trans*-permethrin and *cis*-permethrin respectively) indicated that such pyrethroid might be present in the air even they are semi-volatile organic compounds. More accurately, a study by Bomann et al. [42] from the Bayer company measured a mean concentration of cyfluthrin (a pyrethroid with a molecular structure close to the permethrin) of 0.000021 mg/m^3^ at 1m away from a treated net. Such concentration corresponds to 3.46x10^9^ molecules/cm^3^. *Angioy* et al [36] found that only 6 molecules of a pheromone entered in contact with the olfactory sensillum of moth species may induce a physiological response. We therefore hypothesize that mosquito may detect very low concentration of pyrethroid in the air.

Moreover, some nasal trouble (i.e runny nose) have been recorded when LNs were used for the first time [37]. Such observations reinforce the hypothesis that LNs emit chemicals in the air. Regardless these chemicals are insecticide itself, additive chemicals (i.e. fragrances), degradation products, that composed the net, they may be detected by mosquitoes and elicit behavioral modulation.

The behavior of insects, such as mosquitoes, is driven by the perception and integration of odorant signals in antennae and higher brain center. In our study, we observed that *kdr* resistant mosquitoes were more attracted by host odors emanating behind a permethrin-treated net than host odors behind an untreated net (Fig 3C), it indicates that they perceived at distance a difference between ITN and UTN and behaved differently in response. We then hypothesize that mosquitoes are able to detect chemicals released by net with olfactory receptors (Ors) tuned to respond to these chemicals. As an example, authors recently identified one olfactory receptor activated and another inhibited by synthetic pyrethroïd in *Aedes aegypti* [38], suggesting that such OR may also exist in *Anopheles* mosquitoes. The major research perspective raised by our results is therefore to study the chemical and behavioral ecology relative to vector control tools already widespread in endemic country.

## Insecticide resistance & host seeking behavior

Our results also clearly indicated that the *kdr* mutation, or closely linked polymorphisms [39], modulated the host choice of *An. gambiae* mosquitoes in the presence of a ITN. The strains used in our study share the same genetic background. Except if genes coding for ORs sensitive to LN-related odorants are located in flanking region of *Kdr* mutation,, only a pleiotropic effect of the *kdr* mutation affecting the transmission or integration of the neuronal signal in homozygous mosquitoes could explain the different behaviors between genotypes. The *kdr* mutation may influence the transmission of an odorant signal towards higher brain regions by enhancing the closed-state inactivation of the voltage-gated sodium channel, which plays a central role in message propagation in the nervous system. As a consequence, a reduction of neuronal excitability could be observed in *kdr* mutants in comparison to susceptible individuals [40]. All chemical signals are transduced by spike frequencies in the olfactory sensory neurons [41] and the information sent by stimulated or inhibited neurons is treated in the antennal lobe [42]. Therefore, if the neuronal excitability differs in homozygous *kdr* genotypes, the response pattern of the olfactory neurons and subsequently the signal integration and processing in the central nervous system could be altered, leading to a modified motor response, in this case, a difference in host choice.

The present study suggests the existence of interactions between the physiological mechanisms that allow mosquitoes to survive a contact with insecticide and the behavioral response to olfactory cues. These interactions may have major implications in malaria control. As an example, chemicals emanating from the ITNs are strongly related to the presence of human beings. Should it be integrated as an attractive cue for resistant mosquitoes? This may dramatically affects the personal and community protection given by the massive use of ITNs. Our study only focused on the Kdr mutation, but the resistance pattern in wild *Anophele*s populations is far more complex [43]. It would be interesting to investigate the interaction between each resistance mechanisms isolated in specific strains before going to study this interaction between resistance, behavior and ITNs in semi-field and natural conditions. Recent papers were modeling and questioning the risk conferred by resistance, based on survival to insecticide exposure [44], but the impact of such resistance on behavior is also to be investigated urgently [45].

We used rabbit as an odor source because mosquitoes were fed on rabbits at the laboratory, and were likely “;selected” to respond to rabbit’s odor. But in the field, *Anopheles gambiae* prefers to bite human when available [46]. Whether the same experiment conducted with humans as an odor source will provide similar results remain to be experimentally evaluated. If we used a human instead of the rabbit we change the composition of odor plume (quantity and quality of semiochemicals). Therefore the interaction between chemicals released by LLIN and human odor should induce a different behavior. Nevertheless, our experiment highlighted the involvement of LLIN in host seeking behavior and emphasized the need to studying the relation between LLIN, host odors and mosquito host seeking behavior.

## Kdr genotypes & behavior

Heterozygous RS mosquitoes were more active than SS and RR mosquitoes. The addition of an attractant did not change the proportion of RS leaving the RZ, suggesting that this behavior might be related to a better anemotactic response (i.e response to air flow) or spontaneous flight activity than a better perception of odorants in RS mosquitoes. This hypothesis is strengthened by the absence of difference in the progression towards the olfactometer arms among genotypes. In other words, heterozygous mosquitoes fly more, but might not smell better. On the one hand, by flying more they might increase the probability of encountering a host odorant plume which might be advantageous. Such heterozygous advantage for the *kdr* locus in *An. gambiae s.s.* has been recently documented also for another behavioral trait: the ability to find a hole in a piece of bed net [24] and for male mating [47]. In other hand, it could represent a cost for mosquitoes if energy spent during flight is no more available for other traits closely related to fitness as fertility, fecundity and longevity. This trade off must be deeply investigated as this might have great influence on *Plasmodium* transmission.

The behavior of *kdr* heterozygous individuals in our study must be interpreted with caution because other loci, distinct from the *kdr* locus, could also influence this behavioral trait. Introgression and selection the *kdr* allele to produce the homozygous resistant strain was indeed likely to also have selected linked polymorphisms [45].

## Conclusion

In conclusion, our study showed that the *Anopheles* mosquitoes detected the presence of both physical and chemical barriers of ITNS. Face to this results, it urges to decipher with the interaction between host-seeking behavior, insecticide resistance and vector control tools. The most overlooked part of the puzzle is the chemical ecology in a context of large vector control measure deployment. This research avenue will be challenging for the vector control community but is crucial not to waste forces in wrong directions.

## Acknowledgement

We thank Teun Dekker for helpful discussion. We would like to thank the IEMTV staff in Abomey Calavi (Benin) for technical assistance.

## Supporting information

**S1: Effect of treatment on environment variables**

## References

[1] Cohuet A, Harris C, Robert V, Fontenille D. Evolutionary forces on Anopheles: what makes a malaria vector? Trends Parasitol 2010;26:130–6. doi:10.1016/j.pt.2009.12.001.

[2] Garrett-Jones C, Shidrawi GR. Malaria vectorial capacity of a population of Anopheles gambiae: an exercise in epidemiological entomology. Bull World Health Organ 1969;40.

[3] Garrett-Jones C, Boreham PFL, Pant CP. Feeding habits of anophelines (Diptera: Culicidae) in 1971–78, with reference to the human blood index: a review. Bull Entomol Res 1980;70:165–85.

[4] Coluzzi M, Sabatini A, Petrarca V, Deco MAD. Chromosomal differentiation and adaptation to human environments in the Anopheles gambiae complex. Trans R Soc Trop Med Hyg 1979;73:483–97. doi:10.1016/0035-9203(79)90036-1.

[5] Zaim M, Aitio A, Nakashima N. Safety of pyrethroid-treated mosquito nets. Med Vet Entomol 2000;14:1–5.

[6] Ranson H, N’Guessan R, Lines J, Moiroux N, Nkuni Z, Corbel V. Pyrethroid resistance in African anopheline mosquitoes: what are the implications for malaria control? Trends Parasitol 2011;27:91–8.

[7] Ranson H, Jensen B, Vulule J, Wang X, Hemingway J, Collins F. Identification of a point mutation in the voltage-gated sodium channel gene of Kenyan Anopheles gambiae associated with resistance to DDT and pyrethroids. Insect Mol Biol 2001;9:491–7.

[8] Gatton ML, Chitnis N, Churcher T, Donnelly MJ, Ghani AC, Godfray HCJ, et al. The Importance of Mosquito Behavioural Adaptations to Malaria Control in Africa. Evolution 2013;67:1218–30. doi:10.1111/evo.12063.

[9] Georghiou GP. The evolution of resistance to pesticides. Annu Rev Ecol Syst 1972;3:133–68.

[10] Corbel V, Akogbeto M, Damien GB, Djenontin A, Chandre F, Rogier C, et al. Combination of malaria vector control interventions in pyrethroid resistance area in Benin: a cluster randomized controlled trial. Lancet Infect Dis 2012.

[11] Moiroux N, Gomez MB, Pennetier C, Elanga E, Djènontin A, Chandre F, et al. Changes in Anopheles funestus Biting Behavior Following Universal Coverage of Long-Lasting Insecticidal Nets in Benin. J Infect Dis 2012.

[12] Ndiath MO, Mazenot C, Sokhna C, Trape J-F. How the Malaria Vector Anopheles gambiae Adapts to the Use of Insecticide-Treated Nets by African Populations. PloS One 2014;9.

[13] Lockwood JA, Sparks TC, Story RN. Evolution of Insect Resistance to Insecticides: A Reevaluation of the Roles of Physiology and Behavior. Bull ESA 1984;30:41–51.

[14] Chareonviriyaphap T, Bangs MJ, Suwonkerd W, Kongmee M, Corbel V, Ngoen-Klan R. Review of insecticide resistance and behavioral avoidance of vectors of human diseases in Thailand. Parasit Vectors 2013;6:280. doi:10.1186/1756-3305-6-280.

[15] Miller JR, Siegert PY, Amimo FA, Walker ED. Designation of Chemicals in Terms of the Locomotor Responses They Elicit From Insects: An Update of Dethier et al. (1960). J Econ Entomol 2009;102:2056–60. doi:10.1603/029.102.0606.

[16] Chareonviriyaphap T, Roberts D, Andre RG, Harlan HJ, Manguin S, Bangs MJ. Pesticide avoidance behavior in Anopheles albimanus, a malaria vector in the Americas. J Am Mosq Control Assoc-Mosq News 1997;13:171–83.

[17] Malima R, Oxborough R, Tungu P, Maxwell C, Lyimo I, Mwingira V, et al. Behavioural and insecticidal effects of organophosphate, carbamate and pyrethroid treated mosquito nets against African malaria vectors. Med Vet Entomol 2009;23:317–25.

[18] Achee NL, Sardelis MR, Dusfour I, Chauhan KR, Grieco JP. Characterization of Spatial Repellent, Contact Irritant, and Toxicant Chemical Actions of Standard Vector Control Compounds 1. J Am Mosq Control Assoc 2009;25:156–67.

[19] Deletre E, Martin T, Campagne P, Bourguet D, Cadin A, Menut C, et al. Repellent, Irritant and Toxic Effects of 20 Plant Extracts on Adults of the Malaria Vector Anopheles gambiae Mosquito. PLoS ONE 2013;8:e82103. doi:10.1371/journal.pone.0082103.

[20] Siegert PY, Walker E, Miller JR. Differential behavioral responses of Anopheles gambiae (Diptera: Culicidae) modulate mortality caused by pyrethroid-treated bednets. J Econ Entomol 2009;102:2061–71.

[21] Corbel V, Chabi J, Dabire RK, Etang J, Nwane P, Pigeon O, et al. Field efficacy of a new mosaic long-lasting mosquito net (PermaNet 3.0) against pyrethroid-resistant malaria vectors: a multi centre study in Western and Central Africa. Malar J 2010;9. doi:10.1186/1475-2875-9-113.

[22] Lindsay SW, Adiamah JH, Miller JE, Armstrong JRM. Pyrethroid-treated bednet effects on mosquitoes of the Anopheles gambiae complex in The Gambia. Med Vet Entomol 1991;5:477–83. doi:10.1111/j.1365-2915.1991.tb00576.x.

[23] Pleass R j., Armstrong J r. m., Curtis C f., Jawara M, Lindsay S w. Comparison of permethrin treatments for bednets in The Gambia. Bull Entomol Res 1993;83:133–9. doi:10.1017/S0007485300041870.

[24] Diop MM, Moiroux N, Chandre F, Martin-Herrou H, Milesi P, Boussari O, et al. Behavioral Cost & Overdominance in Anopheles gambiae. PLoS ONE 2015;10:e0121755. doi:10.1371/journal.pone.0121755.

[25] Parker JEA, Angarita-Jaimes N, Abe M, Towers CE, Towers D, McCall PJ. Infrared video tracking of Anopheles gambiae at insecticide-treated bed nets reveals rapid decisive impact after brief localised net contact. Sci Rep 2015;5:13392. doi:10.1038/srep13392.

[26] Grieco JP, Achee NL, Chareonviriyaphap T, Suwonkerd W, Chauhan K, Sardelis MR, et al. A New Classification System for the Actions of IRS Chemicals Traditionally Used For Malaria Control. PLOS ONE 2007;2:e716. doi:10.1371/journal.pone.0000716.

[27] Martinez-Torres D, Chandre F, Williamson M, Darriet F, Berge JB, Devonshire AL, et al. Molecular characterization of pyrethroid knockdown resistance (kdr) in the major malaria vector Anopheles gambiae ss. Insect Mol Biol 1998;7:179–84.

[28] Djegbé I. Modification physiologique et comportementale induites par la resistancé aux insecticides chez les vecteurs du paludisme au Bénin. Université d’Abomey Calavi, 2013.

[29] N’Guessan R, Corbel V, Akogbeto M, Rowland M. Reduced efficacy of insecticide-treated nets and indoor residual spraying for malaria control in pyrethroid resistance area, Benin. Emergent Infect Dis 2007;13:199–206.

[30] Geier M, Boeckh J. A new Y-tube olfactometer for mosquitoes to measure the attractiveness of host odours. Entomol Exp Appl 1999;92:9–19.

[31] R Core Team. R: A language and environment for statistical computing. Vienna, Austria: R Foundation for Statistical Computing; 2014.

[32] Bates D, Maechler M, Bolker B, Walker S. lme4: Linear mixed-effects models using Eigen and S4. R package version 1.1-7 2014.

[33] Takken W, Knols BGJ, Otten H. Interactions between physical and olfactory cues in the host seeking behaviour of mosquitoes: the role of relative humidity. Ann Trop Med Parasitol 1997;91:119–20. doi:10.1080/00034989761427.

[34] Dekker T, Takken W, Carde RT. Structure of host-odour plumes influences catch of Anopheles gambiae ss and Aedes aegypti in a dual-choice olfactometer. Physiol Entomol 2001;26:124–34.

[35] Cardé RT, Gibson G. Host finding by female mosquitoes: mechanisms of orientation to host odours and other cues. Olfaction Vector-Host Interact., Willem Takken and Bart G.J. Knols. Wageningen Academic Publishers; 2010.

[36] Angioy AM, Desogus A, Barbarossa IT, Anderson P, Hansson BS. Extreme sensitivity in an olfactory system. Chem Senses 2003;28:279–84.

[37] WHO. Guidelines for laboratory and field-testing of long-lasting insecticidal nets. 2013.

[38] Bohbot JD, Fu L, Le TC, Chauhan KR, Cantrell CL, Dickens JC. Multiple activities of insect repellents on odorant receptors in mosquitoes. Med Vet Entomol 2011;25:436–44. doi:10.1111/j.1365-2915.2011.00949.x.

[39] Mitri C, Markianos K, Guelbeogo WM, Bischoff E, Gneme A, Eiglmeier K, et al. The kdr508 bearing haplotype and susceptibility to Plasmodium falciparum in Anopheles gambiae: genetic correlation and functional testing. Malar J 2015;14. doi:10.1186/s12936-015-0924-8.

[40] Vais H, Williamson MS, Goodson SJ, Devonshire AL, Warmke JW, Usherwood PN, et al. Activation of Drosophila sodium channels promotes modification by deltamethrin. Reductions in affinity caused by knock-down resistance mutations. J Gen Physiol 2000;115:305–18.

[41] Guidobaldi F, May-Concha IJ, Guerenstein PG. Morphology and physiology of the olfactory system of blood-feeding insects. J Physiol Paris 2014. doi:10.1016/j.jphysparis.2014.04.006.

[42] Hanson B, Christensen TA. Functional characteristics of the antennal lobe. Insect Olfaction. Springer, Berlin: B.S. Hansson; 1999, p. 126–57.

[43] Corbel V, N’guessan R, Brengues C, Chandre F, Djogbenou L, Martin T, et al. Multiple insecticide resistance mechanisms in Anopheles gambiae and Culex quinquefasciatus from Benin, West Africa. Acta Trop 2007;101:207–16.

[44] Viana M, Hughes A, Matthiopoulos J, Ranson H, Ferguson HM. Delayed mortality effects cut the malaria transmission potential of insecticide-resistant mosquitoes. Proc Natl Acad Sci 2016;113:8975–80. doi:10.1073/pnas.1603431113.

[45] Thomas MB, Read AF. The threat (or not) of insecticide resistance for malaria control. Proc Natl Acad Sci 2016;113:8900–2. doi:10.1073/pnas.1609889113.

[46] Lefèvre T, Gouagna L-C, Dabiré KR, Elguero E, Fontenille D, Renaud F, et al. Beyond nature and nurture: phenotypic plasticity in blood-feeding behavior of Anopheles gambiae ss when humans are not readily accessible. Am J Trop Med Hyg 2009;81.

[47] Platt N, Kwiatkowska RM, Irving H, Diabaté A, Dabire R, Wondji CS. Target-site resistance mutations (kdr and RDL), but not metabolic resistance, negatively impact male mating competiveness in the malaria vector Anopheles gambiae. Heredity 2015;115:243–52. doi:10.1038/hdy.2015.33.

[48] Berticat C, Boquien G, Raymond M, Chevillon C. Insecticide resistance genes induce a mating competition cost in Culex pipiens mosquitoes. Genet Res 2002;79:41–7.

